## Supplementary figures and images for "Dynamics of transcriptional programs and chromatin accessibility in mouse spermatogonial cells from early postnatal to adult life"

### Suppl Fig 1

#### **Figure S1.** Lazar-Contes et al.

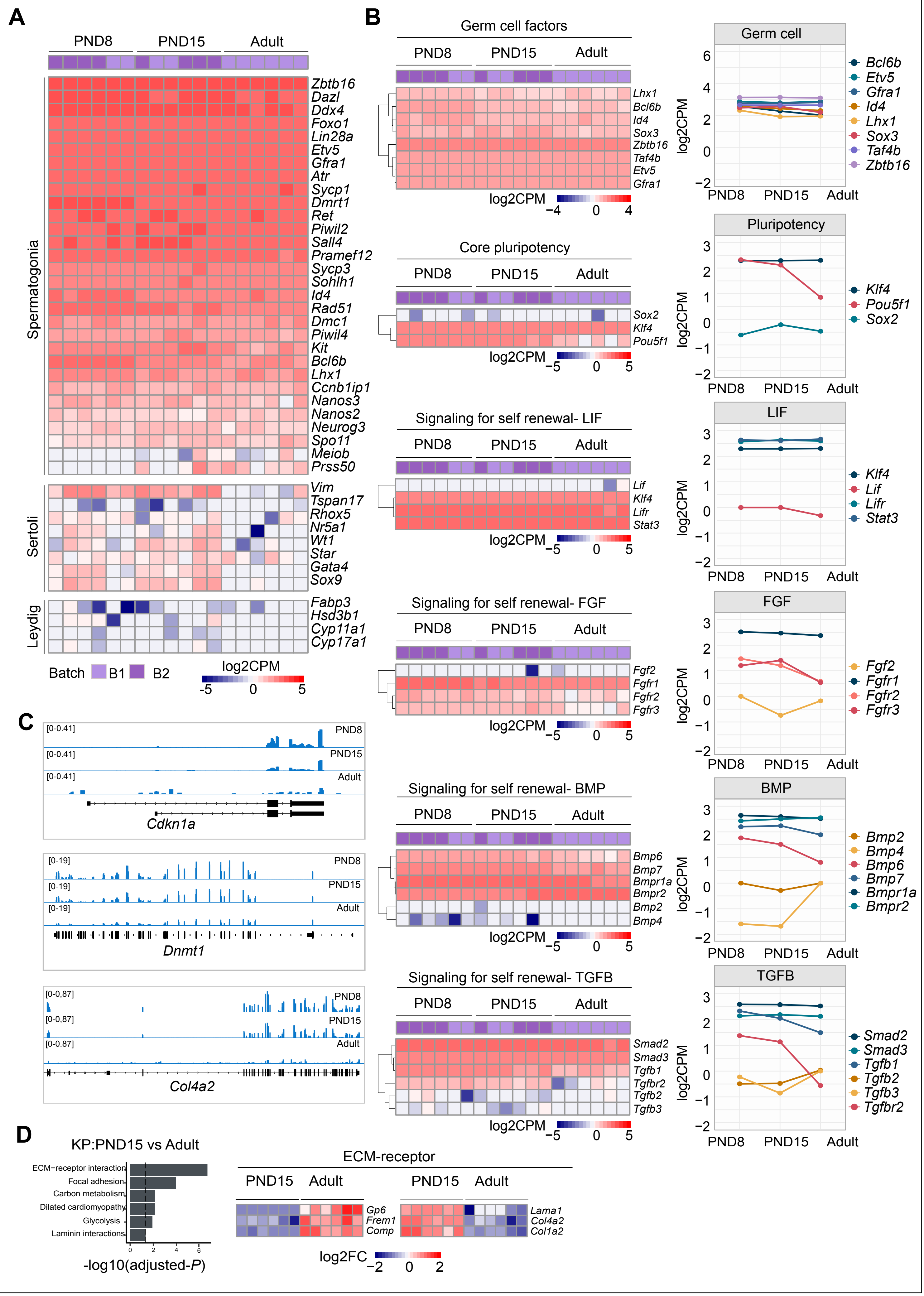

### Suppl Fig 2

#### Figure S2. Lazar-Contes et al.

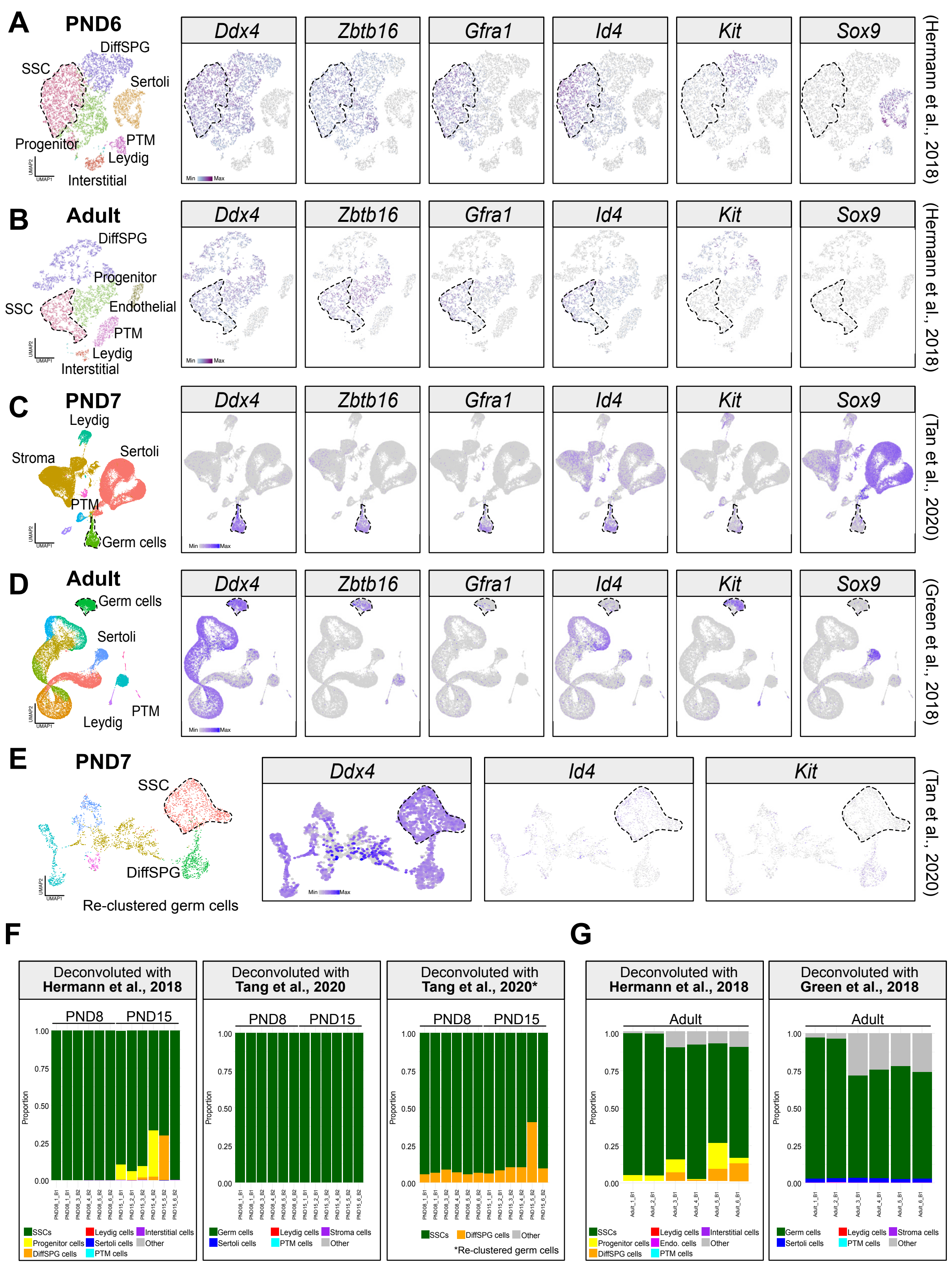

### Suppl Fig 3

**Figure S3.** Lazar-Contes et al.

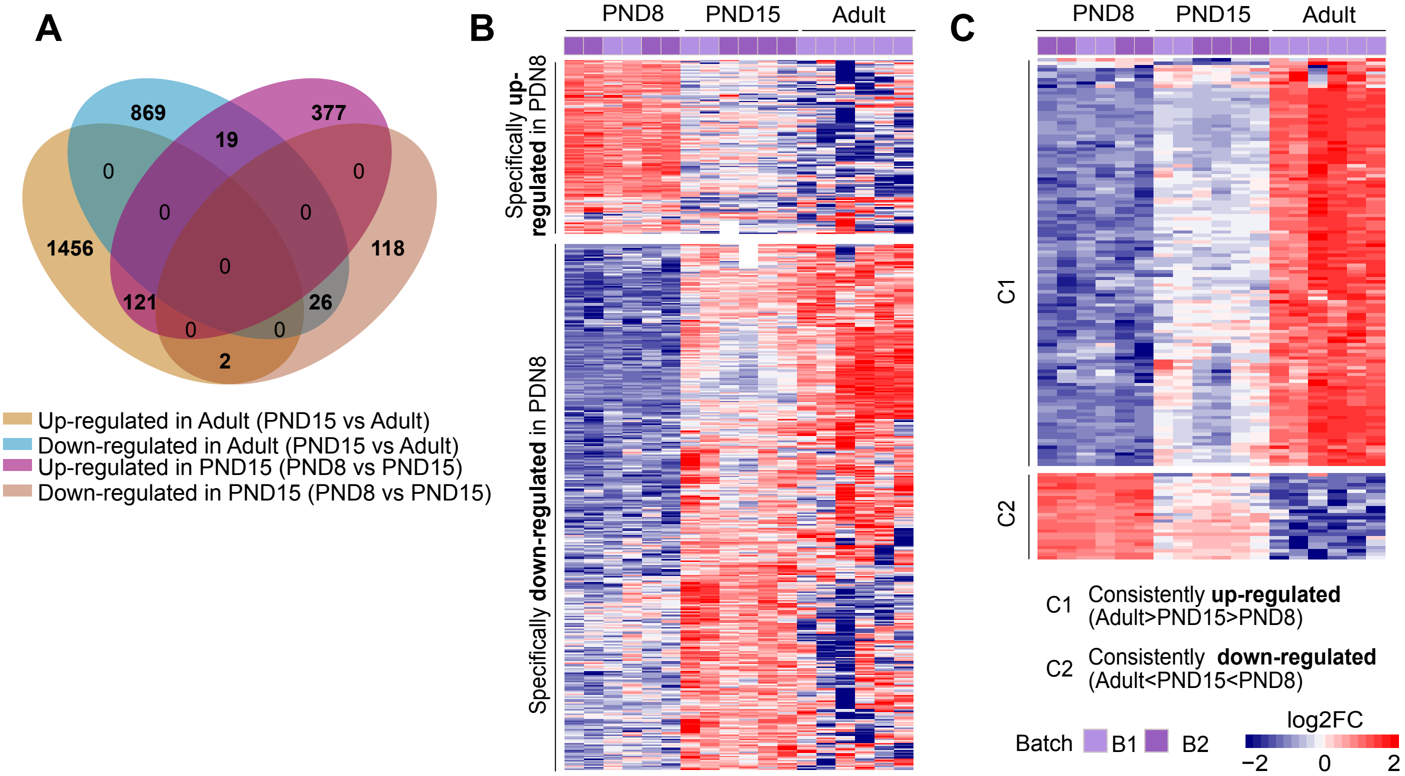

### Suppl Fig 4

**Figure S4.** Lazar-Contes et al.

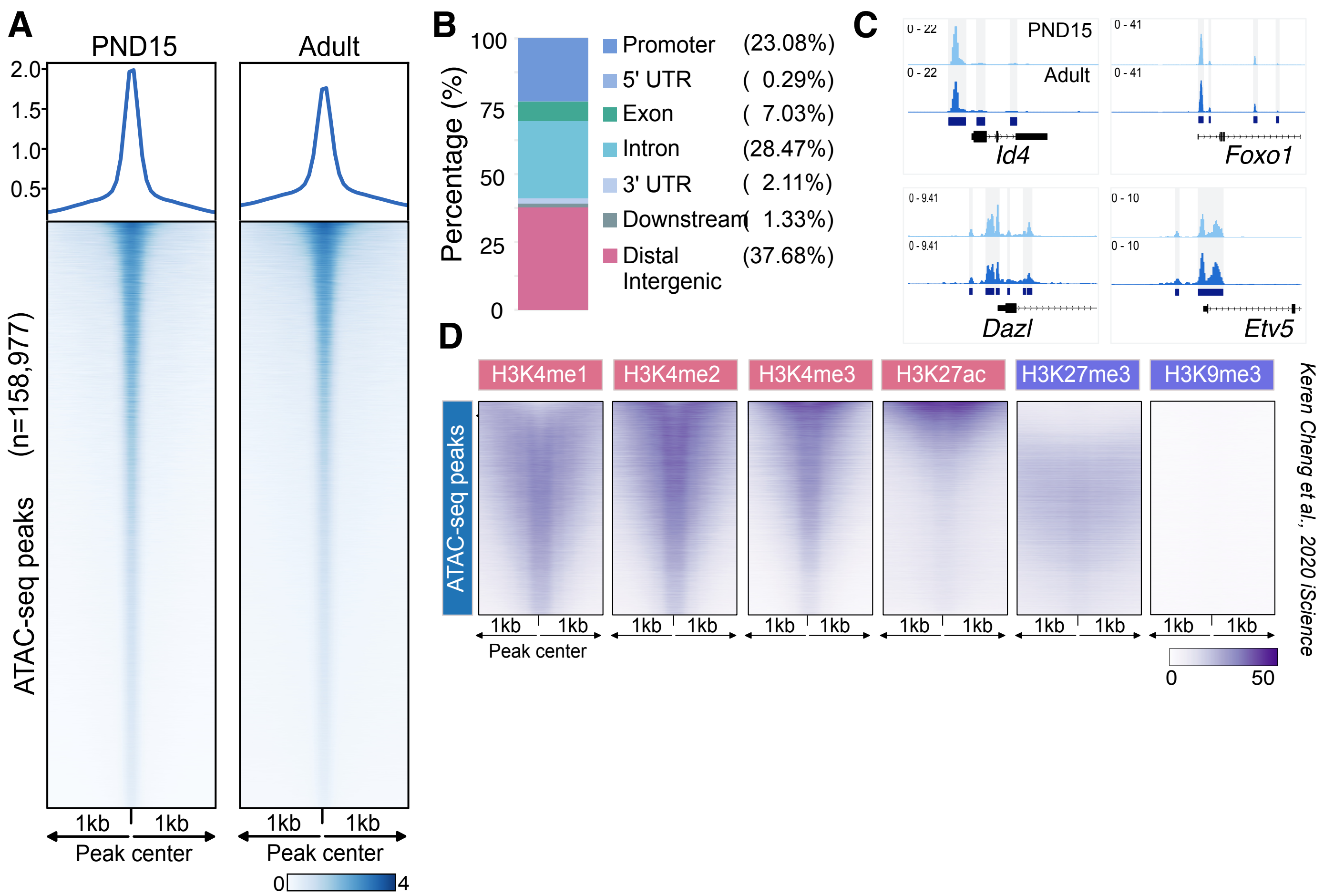
